# Novel retinoic acid receptor-related orphan receptor gamma inverse agonist enhances checkpoint blockade in hepatocellular carcinoma: a preclinical mouse study

**DOI:** 10.64898/2026.10.13.758999

**Authors:** Sabrina Leo, Iqraa Dhoparee-Doomah, Sarita Negi, Ahmed Fouda, France Bourdeau, Gertruda Evaristo, Steven Paraskevas, Jonathan Cools-Lartigue, Jean Tchervenkov

## Abstract

**Background and Aims:** Neutrophil-associated inflammation is associated with poor outcomes in hepatocellular carcinoma (HCC), but tumour-intrinsic regulators of neutrophil recruitment and checkpoint response remain unclear. We evaluated whether inhibiting retinoic acid receptor-related orphan receptor gamma (RORγ) alters neutrophil recruitment, programmed death-ligand 1 (PD-L1) expression, and response to PD-L1 blockade.

**Approach and Results:** Survival associations were analysed in The Cancer Genome Atlas HCC cohort. Neutrophil migration toward HCC cells and PD-L1 expression were assessed after *RORC* silencing or TF-S10 treatment. Immunocompetent mice bearing orthotopic Hep-55.1c tumours received vehicle, TF-S10, anti-PD-L1, or combination treatment. Higher *S100A9* and *CXCL8* expression was associated with shorter survival (*P* = 0.000459 and *P* = 0.0004413, respectively), and tumour-bearing mice had higher neutrophil-to-lymphocyte ratios (*P* < 0.0001). RORγ targeting reduced neutrophil migration and intratumoural Ly6G⁺ infiltration while increasing tumour-cell PD-L1 expression in vitro. Combination treatment reduced endpoint liver tumour volume versus anti-PD-L1 monotherapy (*P* < 0.01). Half of combination-treated mice had tumour volumes ≤0.10 cm³ and none exceeded 1.00 cm³, compared with 62% of controls exceeding 1.00 cm³. Histologically, 85% of combination-treated mice had ≤10% liver tumour involvement, versus no controls. Combination treatment also reduced tumour-associated neutrophil populations and peripheral neutrophil-to-lymphocyte ratios. All active treatments reduced pulmonary metastatic burden versus controls, without significant differences among treatment groups; 53% of combination-treated mice had no detectable lung metastases.

**Conclusions:** RORγ inhibition reduces neutrophil recruitment and increases tumour-cell PD-L1 expression. Combined RORγ inhibition and PD-L1 blockade improves hepatic tumour control in preclinical HCC and warrants further investigation.

## INTRODUCTION

Hepatocellular carcinoma (HCC) is the most common primary liver malignancy and remains a leading cause of cancer-related mortality worldwide^1^. ICIs targeting the programmed cell death protein 1 and programmed death ligand 1 (PD-1/PD-L1) axis have transformed the systemic treatment of advanced HCC, particularly when combined with anti-angiogenic agents or cytotoxic T-lymphocyte-associated protein 4 (CTLA-4) blockade, and ICI-based combinations are now established first-line treatment options for advanced or unresectable HCC. Nevertheless, objective responses are achieved in only approximately 20-30% of patients, and both primary and acquired resistance remain major therapeutic challenges^2,3^. These limitations have intensified efforts to identify tumour-intrinsic and microenvironmental mechanisms that influence immune evasion and determine responsiveness to checkpoint blockade^4^. The HCC tumour microenvironment is particularly complex, arising within a chronically inflamed and frequently fibrotic liver in which malignant, stromal and immune-cell compartments engage in reciprocal signaling that can promote tumour progression and suppress anti-tumour immunity^5^.

Among the immune populations present in HCC, neutrophils have emerged as important regulators of tumour progression and therapeutic resistance. Tumour-associated neutrophils (TANs) are highly plastic and may acquire tumour-promoting functions in response to local cytokines, chemokines and metabolic signals. Increased neutrophil infiltration has been associated with aggressive clinicopathological features, earlier recurrence and reduced survival in HCC^6^. Consistent with this, elevated pretreatment neutrophil-to-lymphocyte ratio (NLR) has been associated with poorer outcomes in patients with HCC receiving ICIs^7^. Mechanistically, TANs can sustain inflammation, promote angiogenesis and extracellular-matrix remodelling, facilitate invasion and metastatic dissemination, and suppress anti-tumour T cell responses through reactive oxygen species (ROS), arginase, proteases and other immunoregulatory mediators^6^. Tumour-derived CXCR2 ligands, including CXCL5, can establish chemotactic gradients that recruit neutrophils into HCC lesions, thereby promoting primary tumour growth and pulmonary metastasis^8^. Furthermore, TANs can recruit macrophages and regulatory T (Treg) cells, reinforcing an immunosuppressive microenvironment and contributing to resistance to sorafenib^9^. These findings suggest that disrupting tumour-driven neutrophil recruitment may improve tumour control and potentially alleviate barriers to effective immunotherapy.

Retinoic acid receptor-related orphan receptor gamma (RORγ), encoded by *RORC*, is a ligand-regulated nuclear receptor that exists as the broadly expressed RORγ isoform and the immune-specific RORγt isoform. RORγt is a lineage-defining transcription factor for interleukin 17-producing lymphocytes and supports inflammatory cytokine programs capable of indirectly promoting granulopoiesis and neutrophil recruitment^10^. In cancer, *RORC* has also been implicated in tumour-associated emergency granulo-monocytopoiesis and the persistence of immunosuppressive myeloid populations, which include large populations of neutrophils, indicating that RORγ-associated signaling can influence systemic myeloid responses^11^. Beyond these immune functions, emerging evidence demonstrates that RORγ can act within malignant cells. In HCC, RORγ has been reported to activate a matrisome-associated transcriptional program that promotes extracellular-matrix deposition, tumour growth and metastatic progression^12^. Pharmacological RORγ antagonism reduced tumour growth and metastasis in preclinical HCC models and enhanced sensitivity to sorafenib^12^; however, whether tumour-cell-intrinsic RORγ regulates immune-checkpoint expression or secreted programs that control neutrophil recruitment in HCC remains unknown.

In this study, we investigated the role of RORγ in HCC tumour-immune crosstalk and evaluated the immune-modulatory activity of TF-S10, a novel RORγ inverse agonist^13,14^. Using genetic and pharmacological approaches, we examined whether RORγ inhibition alters tumour-cell PD-L1 expression and the capacity of HCC cells to recruit neutrophils. We further evaluated the effects of TF-S10 on intratumoural neutrophil accumulation and determined whether RORγ inhibition enhances the anti-tumour efficacy of PD-L1 blockade in an immunocompetent orthotopic HCC model. We demonstrate that RORγ inhibition increases tumour-cell PD-L1 expression while reducing neutrophil migration, intratumoural accumulation and the peripheral neutrophil-to-lymphocyte ratio (NLR). Moreover, combined TF-S10 and anti-PD-L1 therapy produced the greatest reduction in primary liver tumour burden, while all treatment groups reduced pulmonary metastatic burden compared with controls. These findings identify RORγ as a regulator of HCC tumour-immune interactions and provide a preclinical rationale for combining RORγ inhibition with PD-L1 blockade.

## MATERIALS AND METHODS

### Cell culture

Huh-7 cells were obtained from the Transplant-Immunology Laboratory of Drs. Tchervenkov and Paraskevas (RI-MUHC) and cultured in DMEM without sodium pyruvate supplemented with 10% heat-inactivated FBS and 1% penicillin-streptomycin. Hepa1-6 cells (ATCC) were cultured in DMEM with 10% FBS, 1% glutamine and 1% penicillin-streptomycin, whereas Hep-55.1c cells (Meyer Cancer Center) were maintained in RPMI 1640 with 10% FBS, 1× non-essential amino acids, 1 mM sodium pyruvate, 2 mM L-glutamine and 1% penicillin-streptomycin. All cells were maintained at 37°C, 5% CO₂ and routinely screened for mycoplasma.

### Public dataset survival analysis

Overall survival was analyzed using the TCGA liver hepatocellular carcinoma (LIHC) cohort accessed through the UCSC Xena Browser. *S100A9* and *CXCL8* expression data were obtained from RNA-sequencing datasets and reported as fragments per kilobase of transcript per million mapped reads upper quartile (FPKM-UQ). Patients were stratified into high- and low-expression groups according to median expression. Kaplan-Meier survival curves were generated and compared using the log-rank (Mantel-Cox) test.

### siRNA transfection

*RORC* was silenced using small interfering RNA (si*RORC*; Integrated DNA Technologies) delivered with Lipofectamine RNAiMAX (Thermo Fisher Scientific) according to the manufacturer’s instructions. Cells were transfected at approximately 60–70% confluency and maintained for 48 hours before downstream analyses. A non-targeting siRNA (siNC) was used as the negative control. Knockdown efficiency was confirmed by RT–qPCR.

### RT–qPCR

Total RNA was extracted using the RNeasy Mini Kit (Qiagen) according to the manufacturer’s instructions, and RNA concentration and purity were assessed by NanoDrop spectrophotometry. cDNA was synthesized using the High-Capacity cDNA Reverse Transcription Kit (Thermo Fisher Scientific). Quantitative PCR was performed using iTaq Universal SYBR Green Supermix (Bio-Rad) on a CFX Opus 96 Real-Time PCR System (Bio-Rad Laboratories). *CD274* (PD-L1) expression was assessed using gene-specific custom primers (forward: 5′-TGCAGGGCATTCCAGAAAGATG-3′; reverse: 5′-TAGGTCCTTGGGAACCGTGA-3′), with *GAPDH* used as the internal control (forward: 5′-TTGCCCTCAACGACCACTTT-3′; reverse: 5′-TGGTCCAGGGGTCTTACTCC-3′). Relative gene expression was calculated using the 2^−ΔΔCt^ method.

### Neutrophil chemotaxis assay

Huh-7 cells (5 × 10⁴ cells/well) were seeded in the lower chamber of 24-well Transwell plates and after 24 hours, treated with TF-S10 or transfected with si*RORC* (si*RORC*1-3). After 72 hours, the medium was replaced with serum-free DMEM for 24 hours to minimize exogenous chemoattractants. Human neutrophils were then isolated from peripheral blood obtained voluntarily from healthy adult donors, stained with Hoechst dye and added to the upper chamber (5 × 10⁵ cells/insert) in serum-free medium using 3-μm-pore Transwell inserts. After 4 hours, neutrophil migration toward Huh-7 cells was quantified by fluorescence microscopy using an EVOS M7000 Imaging System (Thermo Fisher Scientific), based on the number of cells that migrated to the lower chamber.

### TF-S10 preparation and treatment

TF-S10 was used as a pharmacological inverse agonist of RORγ^13,14^. For *in vitro* experiments, TF-S10 was dissolved in DMSO and diluted in culture medium to the indicated final concentrations, with equivalent DMSO concentrations used as vehicle controls. For *in vivo* experiments, TF-S10 was formulated in 4% Tween-80 in PBS and administered at 25 mg/kg by daily subcutaneous injection. Vehicle-treated mice received 4% Tween-80 in PBS alone.

### Subcutaneous and orthotopic HCC mouse models

All animal experiments were approved by the RI-MUHC Ethics Committee (AUP#8210). Male C57BL/6 mice (7–10 weeks; Charles River Laboratories) were used based on prior model optimization demonstrating more consistent tumour engraftment and growth in males than females. Animals with documented technical failure of tumour-cell injection or implantation were excluded from analysis. For the subcutaneous model, Hep-55.1c cells (1 × 10⁶ in 100 μL PBS) were injected into the left flank. Peripheral blood was analyzed once tumours reached 0.4–0.5 cm³. For the orthotopic model, Hep-55.1c cells (7.5 × 10⁵ in 20 μL PBS) were implanted into the left hepatic lobe under isoflurane anaesthesia. Mice were randomly assigned on day 7 to vehicle/isotype control, TF-S10 (25 mg/kg subcutaneously daily), anti-PD-L1 (10 mg/kg intraperitoneally every 3 days), or combination therapy. At endpoint, hepatic tumours were measured ex vivo by digital calipers and tumour volume calculated using the ellipsoid formula. Animals were euthanized at the scheduled endpoint on day 26.

### Histopathological assessment of liver tumours and pulmonary metastases

At endpoint, livers and lungs were harvested, fixed in 10% neutral-buffered formalin, processed, and H&E stained by the RI-MUHC Immunohistochemistry Facility. Sections were assessed by a trained pathologist (G. Evaristo). Liver tumour burden was quantified as the percentage of liver section occupied by tumour, while pulmonary metastatic burden was assessed by tumour area and number of metastatic foci. Each distinct tumour-cell cluster within the lung parenchyma was counted as one metastatic focus.

### Immunofluorescence staining and quantification of neutrophil infiltration

Orthotopic liver tumours were fixed in 4% paraformaldehyde, cryopreserved, and OCT-embedded. Sections were stained for Ly6G, pan-cytokeratin, and RORγ (PathPlus, LS-A5191), with appropriate fluorescent secondary antibodies and DAPI nuclear counterstain. Images were acquired using a Zeiss LSM880 confocal microscope with a 20x objective under identical settings. Tumour regions were defined by pan-cytokeratin staining, and neutrophil infiltration was quantified as Ly6G⁺ area per field using ImageJ, with thresholds adjusted to exclude nonspecific staining.

### Multiplex immunofluorescence staining and image analysis

Multiplex immunofluorescence staining was performed on formalin-fixed paraffin-embedded mouse tumour sections by the BC Cancer Vancouver laboratory using an OPAL-based protocol. Slides were baked, deparaffinized, re-fixed in 10% neutral-buffered formalin and subjected to heat-induced antigen retrieval. Sequential staining was performed on an Intellipath FLX autostainer using repeated rounds of primary antibody incubation, HRP-polymer detection, OPAL fluorophore amplification and antigen retrieval between rounds. The panel included RORγ (clone EPR20006, Abcam, ab207082; 1:500; OPAL 620), CD163 (clone EPR19518, Abcam, ab182422; 1:2000; OPAL 690), pan-cytokeratin (clone EPR28285-45, Abcam, ab308262; 1:5500; OPAL 570), Ly6G (clone E6Z1T, Cell Signaling Technology, 87048S; 1:100; OPAL 480), PD-L1 (clone F3ZT1T, Cell Signaling Technology, 80517; 1:100; OPAL 520) and CD8α (clone D4W2Z, Cell Signaling Technology, 98941S; 1:100; OPAL 780). Slides were counterstained with DAPI, mounted using ProLong Diamond Antifade Mountant and imaged using a Vectra Polaris system. Image analysis was performed using inForm software.

### Flow cytometry

Peripheral blood was collected from mice, and red blood cells were lysed using BD Pharm Lyse Lysing Buffer (BD Biosciences, 555899) according to the manufacturer’s instructions. Remaining cells were washed with PBS and stained with fluorophore-conjugated surface antibodies for 30 min at room temperature. Surface markers included CD3e PE-Cyanine7, CD11b Brilliant Violet 510, Ly6G FITC, and CD274/PD-L1 APC. For intracellular staining, cells were fixed and permeabilized using the FOXP3 Fixation/Permeabilization Kit (eBioscience) and stained with RORγt PE for 45 min at room temperature.

To assess tumour-cell PD-L1 expression, human Huh-7 and murine Hepa1-6 and Hep-55.1c cells were treated with TF-S10 (5 or 10 μM) or the corresponding controls. Cell-surface PD-L1 expression was assessed by flow cytometry using an APC-conjugated anti-CD274/PD-L1 antibody (clone MIH5; Thermo Fisher Scientific/eBioscience, 17-5982-82). The same antibody was used for human and murine HCC cell lines. PD-L1 expression was quantified as the percentage of PD-L1-positive cells.

Samples were acquired on a BD LSRFortessa flow cytometer and analysed using FlowJo software. Antibody clones, vendors, catalogue numbers, and fluorophores are provided in Supplementary Table 1. The neutrophil-to-lymphocyte ratio (NLR) was calculated as the ratio of CD11b⁺Ly6G⁺ neutrophils to CD3⁺ lymphocytes.

### Statistical analysis

Statistical analyses were performed using GraphPad Prism. Data are presented as mean ± SEM unless otherwise indicated. Two-group comparisons were performed using unpaired two-tailed *t*-tests or Mann–Whitney U tests, as appropriate. Multiple-group comparisons were performed using one-way ANOVA with Šídák’s multiple-comparisons test, Brown–Forsythe and Welch ANOVA followed by Dunnett’s T3 multiple-comparisons test, or Kruskal–Wallis tests with Dunn’s multiple-comparisons test, as appropriate. Kaplan–Meier survival curves were compared using the log-rank (Mantel–Cox) test. A *P* value < 0.05 was considered statistically significant.

## RESULTS

### Neutrophil-associated inflammatory gene expression correlates with poor survival in human HCC, while tumour-bearing mice exhibit increased systemic neutrophil-to-lymphocyte ratio

To establish the clinical relevance of neutrophil-associated inflammation in HCC, we first examined the relationship between the expression of two inflammatory genes, *S100A9* and *CXCL8*, and patient survival using the TCGA-LIHC cohort. *S100A9*, which is highly expressed in neutrophils and other myeloid cells, and *CXCL8*, which encodes a chemokine involved in neutrophil recruitment, were selected to assess distinct aspects of neutrophil-associated inflammation. Kaplan–Meier survival analyses demonstrated that elevated *S100A9* expression was significantly associated with reduced overall survival (Figure 1A; *P* = 0.000459). Similarly, patients with higher *CXCL8* expression exhibited significantly poorer overall survival (Figure 1B; *P* = 0.0004413). These findings support an association between neutrophil-related inflammatory gene expression and adverse clinical outcomes in HCC.

**Fig. 1.**
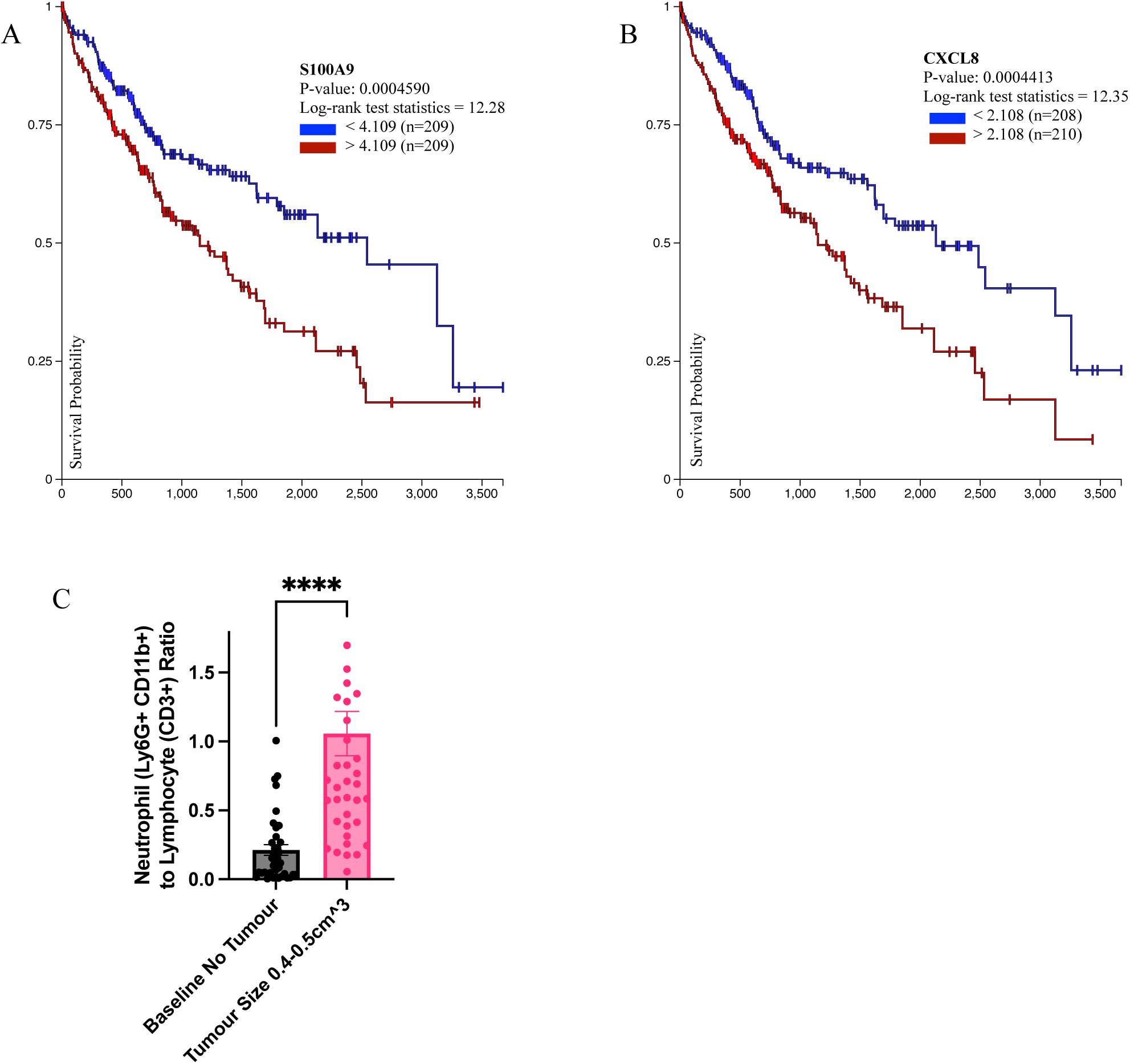
Elevated neutrophil-associated gene expression is associated with poor survival in HCC, while tumour-bearing mice exhibit increased peripheral neutrophil-to-lymphocyte ratios. (A, B) Kaplan–Meier overall survival curves for patients in The Cancer Genome Atlas liver hepatocellular carcinoma (TCGA-LIHC) cohort, stratified by tumour RNA-seq expression (FPKM-UQ) of *S100A9* (A) and *CXCL8* (B). Patients were stratified into low-expression (blue) and high expression (red) groups using the median expression value for each gene, as determined by UCSC Xena Browser. Survival differences between high- and low-expression groups were assessed using the log-rank test (A, *P* = 0.000459; B, *P* = 0.0004413). (C) Peripheral neutrophil-to-lymphocyte ratio, calculated as the ratio of Ly6G⁺CD11b⁺ neutrophils to CD3⁺ lymphocytes, in non-tumour-bearing mice (n = 39) and mice bearing established subcutaneous Hep-55.1c tumours measuring 0.4–0.5 cm³ (n = 40). Data are presented as mean ± SEM. Two-tailed Mann–Whitney U test; ****P < 0.0001.

Given the clinical relevance of neutrophil-associated inflammation, we next sought to characterize systemic neutrophil predominance in our immunocompetent murine HCC model, which was subsequently used to investigate the effects of RORγ inhibition on neutrophil-associated responses. Peripheral blood analysis revealed a significantly elevated neutrophil-to-lymphocyte ratio in mice bearing established subcutaneous Hep-55.1c tumours compared with non-tumour-bearing controls (Figure 1C; *P* < 0.0001). These findings establish the clinical relevance of neutrophil-associated inflammatory markers in human HCC and demonstrate a tumour-associated shift in circulating immune-cell populations in our murine model, providing a rationale for further investigating the role of RORγ in regulating neutrophil recruitment and infiltration.

### RORγ inhibition suppresses neutrophil recruitment in vitro and reduces tumour-associated neutrophil infiltration in vivo

To determine whether tumour-intrinsic RORγ influences neutrophil recruitment, neutrophil chemotaxis toward Huh-7 HCC cells was assessed using an *in vitro* Transwell migration assay. Silencing of *RORC* significantly reduced neutrophil migration compared with siNC controls (Figure 2A), suggesting that RORγ expression in HCC cells contributes to neutrophil-attracting signals. Consistently, pharmacological inhibition of RORγ with TF-S10 (5 and 10 μM) significantly decreased neutrophil migration toward Huh-7 cells compared with DMSO-treated controls (Figure 2B). To determine whether these findings were reflected *in vivo*, Ly6G immunofluorescence was used to quantify neutrophil infiltration in orthotopic Hep-55.1c liver tumours. TF-S10-treated mice exhibited a significant reduction in Ly6G⁺ area compared with controls (Figure 2C), indicating decreased neutrophil accumulation within the tumour microenvironment. Representative immunofluorescence images illustrated reduced Ly6G⁺ neutrophil infiltration in TF-S10-treated tumours (Figure 2D, E). Furthermore, the number of Ly6G⁺RORγ⁺ double-positive cells was significantly reduced following TF-S10 treatment (Figure 2F), indicating a decrease in RORγ-expressing neutrophils within the tumour microenvironment. Collectively, these findings demonstrate that RORγ targeting reduces neutrophil migration toward HCC cells and decreases tumour-associated neutrophil infiltration *in vivo*.

**Fig. 2.**
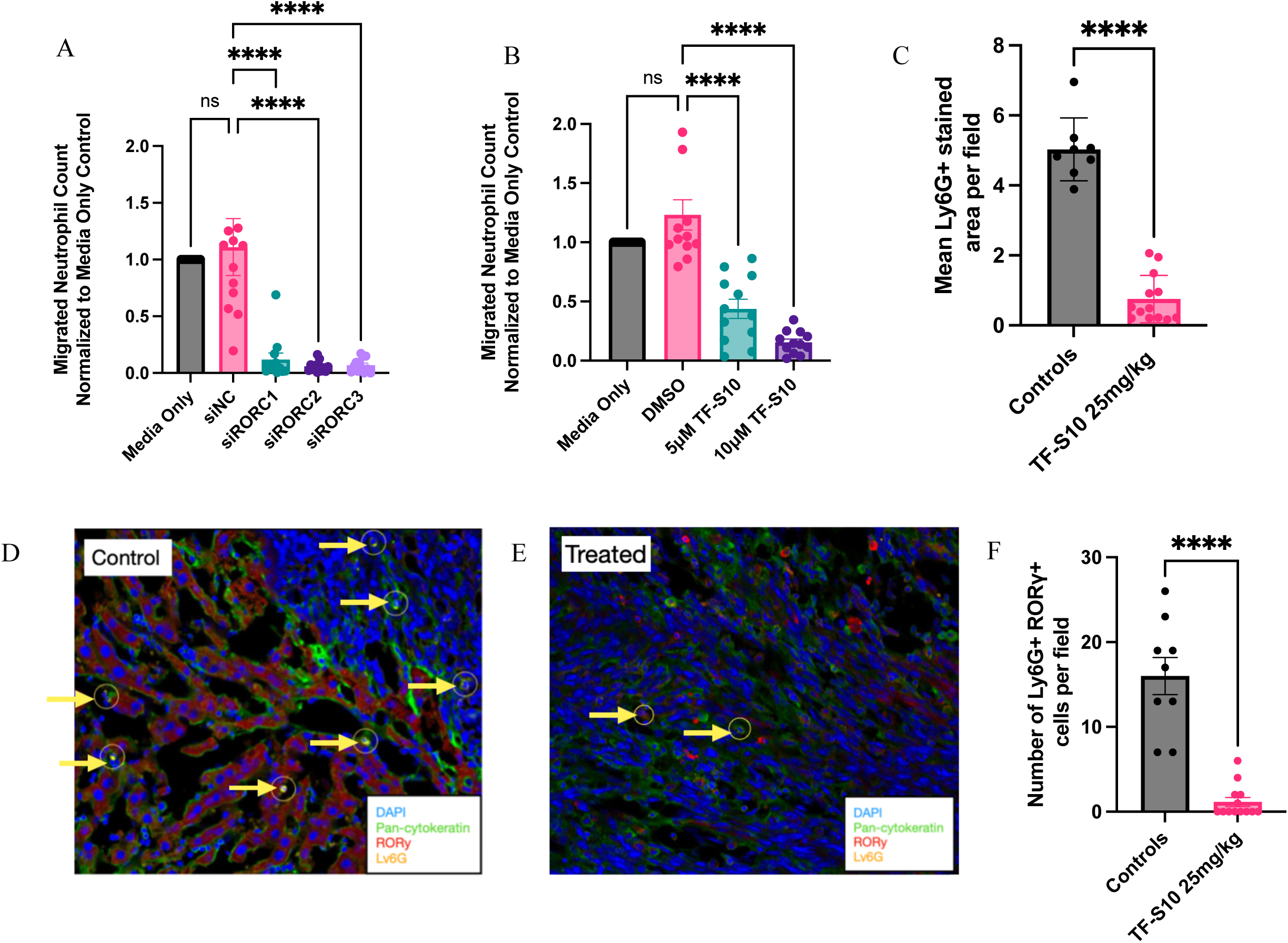
RORγ targeting reduces neutrophil chemotaxis in vitro and tumour-associated neutrophil infiltration in vivo. (A, B) Neutrophil chemotaxis toward Huh-7 cells following *RORC* silencing with siRORC1–3 (A) or treatment with TF-S10 (5 or 10 μM) (B). Migrated neutrophil counts were normalized to the medium-only control. (C) Quantification of the mean area occupied by Ly6G⁺ staining in orthotopic liver tumours sections from control (n = 8) and TF-S10-treated (25 mg/kg; n = 13) mice. (D, E) Representative immunofluorescence images of tumour sections from control (D) and TF-S10-treated (E) mice, acquired using a 20x objective, stained for DAPI (blue), pan-cytokeratin (green), RORγ (red), and Ly6G (yellow). Yellow arrows indicate Ly6G⁺RORγ⁺ cells. (F) Quantification of manually counted Ly6G⁺RORγ⁺ double-positive cells per 20x microscopic field in tumour sections from control (n = 9) and TF-S10-treated (25 mg/kg; n = 13) mice. Data are presented as mean ± SEM. Statistical significance was assessed using one-way ANOVA with Šídák’s multiple-comparisons test (A, B) or a two-tailed Mann–Whitney U test (C, F). ****P < 0.0001; ns, not significant.

### RORγ inhibition increases PD-L1 expression in HCC cells in vitro

To determine whether RORγ regulates immune checkpoint expression in HCC cells, *CD274*/PD-L1 expression was assessed following genetic and pharmacological inhibition of RORγ. Silencing of *RORC* significantly increased *CD274* mRNA expression compared with the non-targeting siNC control (Figure 3A), indicating increased PD-L1 expression at the transcript level. Consistently, treatment with the RORγ inverse agonist TF-S10 (10 μM) significantly increased *CD274* mRNA expression compared with DMSO-treated controls (Figure 3B). Flow cytometric analysis further demonstrated increased PD-L1 surface expression following TF-S10 treatment in human Huh-7 and murine Hep-55.1c cells (Figure 3C, D). A similar increase was observed in Hepa1-6 cells, supporting the reproducibility of this effect across multiple HCC cell lines. Notably, the proportion of PD-L1-positive Hep-55.1c cells approached 100% following TF-S10 treatment (Figure 3D), indicating a marked increase in cell-surface PD-L1 expression. Immunofluorescence analysis further demonstrated a significant increase in PD-L1 integrated fluorescence intensity normalized to cell count following TF-S10 treatment (Figure 3E), with representative images illustrating stronger PD-L1 staining in treated Hep-55.1c cells compared with controls (Figure 3F). Collectively, these findings demonstrate that RORγ inhibition enhances PD-L1 expression at both the transcriptional and protein levels in HCC cells, supporting a role for RORγ in modulating tumour immune checkpoint expression.

**Fig. 3.**
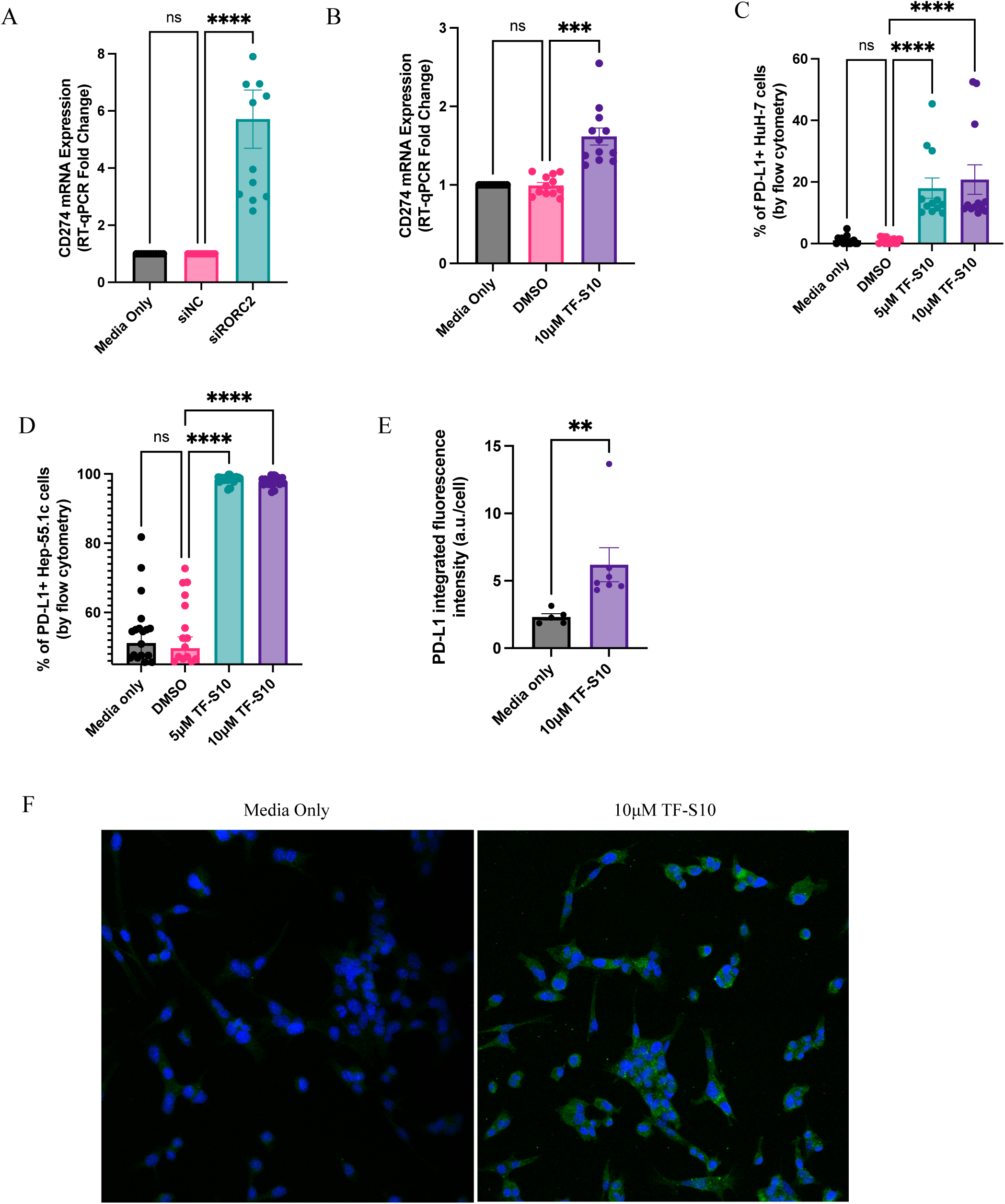
RORγ inhibition enhances PD-L1 expression in HCC cells in vitro. (A, B) *CD274* (PD-L1) mRNA expression following *RORC* silencing with siRORC2 (A) or treatment with TF-S10 (10 μM) (B), assessed by RT-qPCR, in Huh-7 cells. (C, D) Quantification of PD-L1 surface expression by flow cytometry in Huh-7 (C) and Hep-55.1c (D) cells following TF-S10 treatment (5 or 10 μM). (E) Quantification of PD-L1 integrated fluorescence intensity in Hep-55.1c cells following TF-S10 treatment (10 μM), measured from immunofluorescence images using ImageJ software and normalized to cell count (arbitrary units per cell, a.u./cell). (F) Representative immunofluorescence images of Hep-55.1c cells under medium-only and TF-S10-treated (10 μM) conditions, stained for DAPI (blue) and PD-L1 (green). Data are presented as mean ± SEM. Statistical significance was assessed using one-way ANOVA with Šídák’s multiple-comparisons test (A–D) or the Kruskal–Wallis test with Dunn’s multiple-comparisons test (E). **P < 0.01, ***P < 0.001, ****P < 0.0001; ns, not significant.

### Combined RORγ inhibition and anti-PD-L1 therapy reduces gross tumour burden in orthotopic HCC

To determine whether RORγ inhibition could enhance the therapeutic efficacy of immune checkpoint blockade *in vivo*, mice bearing established orthotopic Hep-55.1c liver tumours were treated with TF-S10, anti-PD-L1, or the combination of both therapies (Figure 4A). Endpoint liver tumour volume, measured *ex vivo* using digital calipers following liver harvest, was significantly reduced following TF-S10 monotherapy and combined TF-S10 and anti-PD-L1 treatment compared with controls (Figure 4B). Among the treatment groups, combination therapy resulted in significantly lower tumour volumes than anti-PD-L1 monotherapy (Figure 4C). Analysis of tumour-volume distributions further demonstrated a shift toward lower tumour burden in treated mice, with 50% of combination-treated mice exhibiting minimal tumour burden (≤0.10 cm³) and no mice in this group having tumours exceeding 1.00 cm³ (Figure 4D). Collectively, these findings demonstrate that RORγ inhibition reduces gross tumour burden in orthotopic HCC and that its combination with anti-PD-L1 blockade provides additional tumour control compared with anti-PD-L1 monotherapy.

**Fig. 4.**
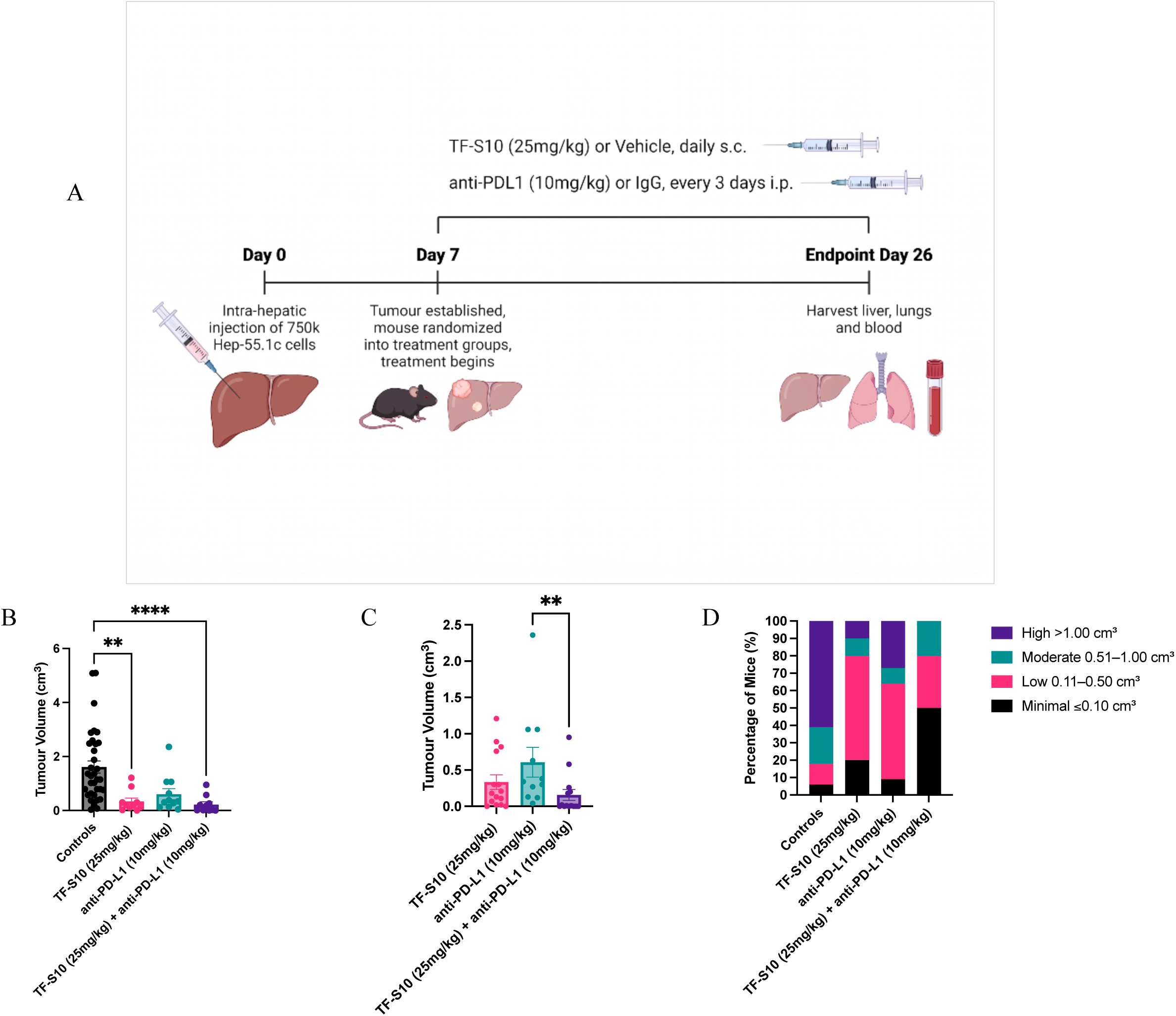
RORγ inhibition combined with anti-PD-L1 blockade reduces gross tumour burden in orthotopic HCC. (A) Schematic of the orthotopic HCC treatment protocol. C57BL/6 mice received intrahepatic injections of 7.5 × 10⁵ Hep-55.1c cells on day 0. On day 7, mice were randomized into four treatment groups: control, TF-S10 (25 mg/kg), anti-PD-L1 (10 mg/kg), or combination therapy. TF-S10 or vehicle was administered daily by subcutaneous injection, while anti-PD-L1 or isotype control IgG was administered intraperitoneally every 3 days. Liver, lungs, and blood were collected on day 26. (B) Endpoint liver tumour volumes measured using calipers across treatment groups (control, n = 43; TF-S10, n = 10; anti-PD-L1, n = 11; combination, n = 10). (C) Comparison of caliper-measured tumour volumes among TF-S10, anti-PD-L1, and combination treatment groups. (D) Distribution of mice according to endpoint tumour volume, categorized as minimal (≤0.10 cm³), low (0.11–0.50 cm³), moderate (0.51–1.00 cm³), or high (>1.00 cm³) tumour burden. Data are presented as mean ± SEM. Statistical significance was assessed using the Kruskal–Wallis test with Dunn’s multiple-comparisons test (B, C). **P < 0.01; ****P < 0.0001.

### Combined RORγ inhibition and anti-PD-L1 therapy reduces histological tumour burden and tumour-cell density in orthotopic HCC

To further evaluate treatment-associated changes in tumour burden, histopathological assessment of hematoxylin and eosin (H&E)-stained liver sections was performed. Quantification of the percentage of liver area occupied by tumour demonstrated significantly reduced histological tumour involvement following TF-S10 monotherapy and combination treatment compared with controls, with the lowest tumour involvement observed in the combination-treated group (Figure 5A). Notably, combination therapy significantly reduced histological tumour burden compared with either monotherapy. Analysis of histological tumour-involvement distributions further demonstrated a shift toward lower tumour burden, with the majority of combination-treated mice exhibiting minimal or no detectable tumour involvement (≤10%), whereas none of the control mice fell within these categories (Figure 5B).

**Fig. 5.**
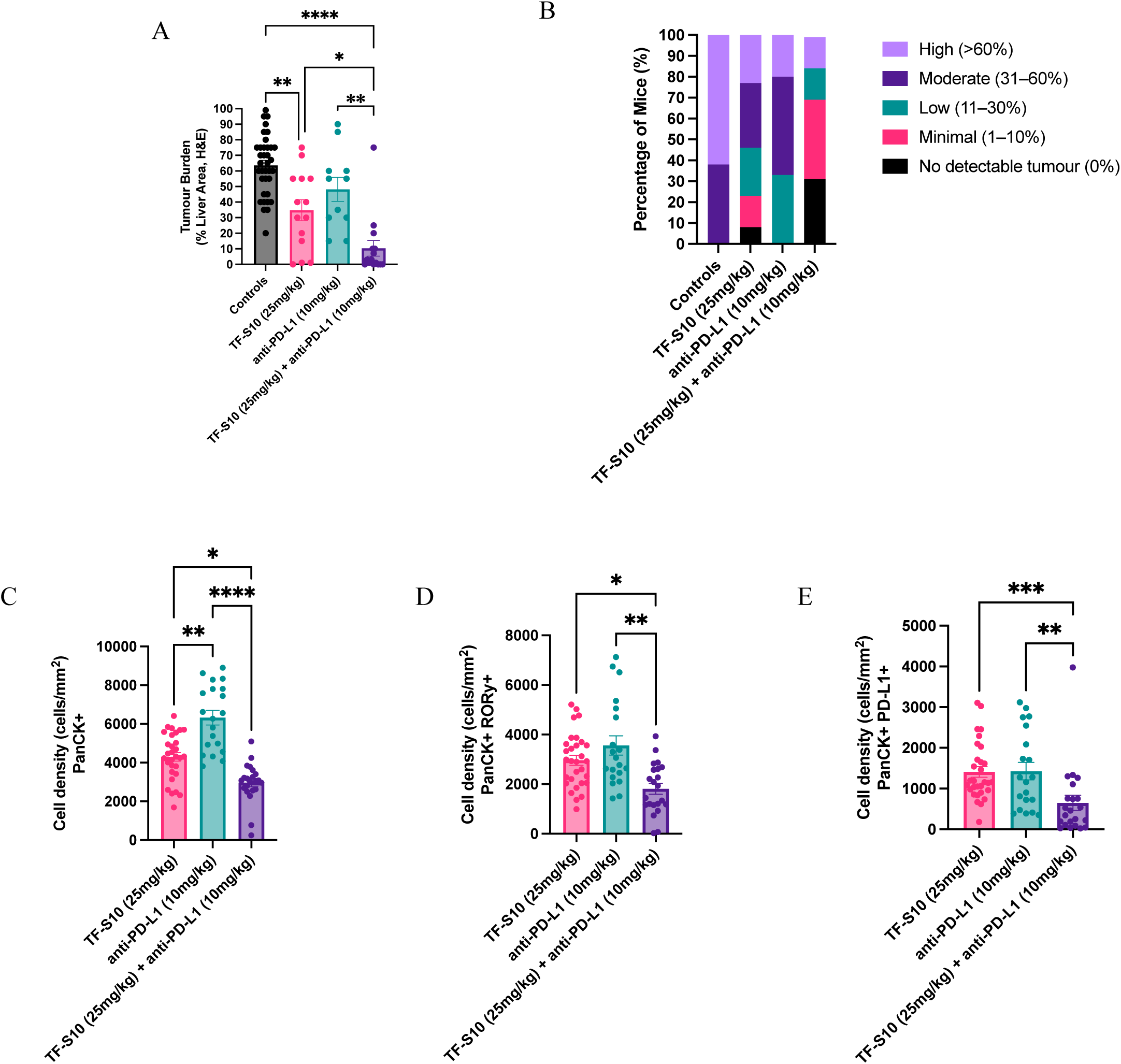
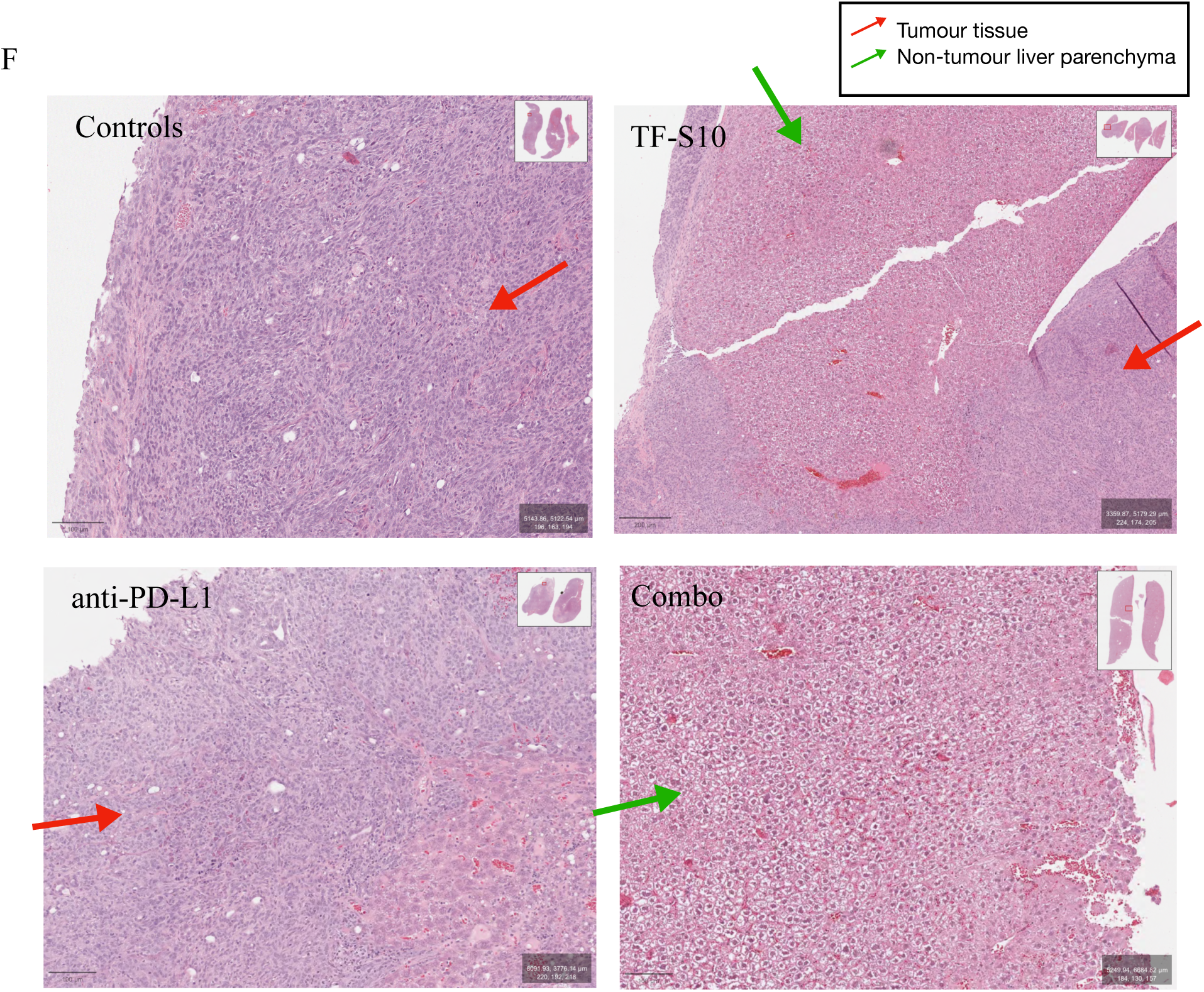
RORγ inhibition combined with anti-PD-L1 blockade reduces histological tumour burden and tumour-cell density in orthotopic HCC. (A) Quantification of histological liver tumour burden, expressed as the percentage of liver area occupied by tumour, assessed using hematoxylin and eosin (H&E)-stained sections from control (n = 36), TF-S10 (25 mg/kg; n = 14), anti-PD-L1 (10 mg/kg; n = 11), and combination (n = 15) treatment groups. (B) Distribution of mice according to histological liver tumour involvement, categorized as no detectable tumour (0%), minimal (1–10%), low (11–30%), moderate (31–60%), or high (>60%) involvement. (C–E) Quantification of pan-cytokeratin-positive (panCK⁺) tumour-cell density (C), panCK⁺RORγ⁺ cell density (D), and panCK⁺PD-L1⁺ cell density (E) in liver tumour sections from TF-S10, anti-PD-L1, and combination treatment groups, expressed as cells/mm². (F) Representative H&E-stained liver sections from control, TF-S10, anti-PD-L1, and combination treatment groups. Red arrows indicate tumour tissue, whereas green arrows indicate non-tumour liver parenchyma. Insets show low-magnification overviews of the corresponding liver sections. Data are presented as mean ± SEM. Statistical significance was assessed using Brown–Forsythe and Welch ANOVA followed by Dunnett’s T3 multiple-comparisons test (A), or the Kruskal–Wallis test with Dunn’s multiple-comparisons test (C–E). *P < 0.05, **P < 0.01, ***P < 0.001, ****P < 0.0001.

Quantitative multiplex immunofluorescence analysis further demonstrated a significant reduction in pan-cytokeratin-positive (panCK⁺) tumour-cell density following combination therapy compared with either monotherapy (Figure 5C), supporting reduced epithelial tumour-cell burden within the liver. Similarly, panCK⁺RORγ⁺ tumour-cell density was significantly decreased in the combination-treated group, indicating a reduction in the density of RORγ-expressing tumour cells (Figure 5D). Furthermore, panCK⁺PD-L1⁺ tumour-cell density was significantly reduced following combination therapy compared with either monotherapy (Figure 5E), reflecting fewer PD-L1-expressing tumour cells within the assessed tissue. Representative H&E-stained liver sections further illustrated differences in tumour involvement across control, TF-S10, anti-PD-L1, and combination-treated groups (Figure 5F). Collectively, these findings demonstrate that combined RORγ inhibition and PD-L1 blockade reduces histological tumour involvement and the density of tumour cells expressing RORγ and PD-L1, providing complementary histopathological evidence of improved tumour control.

### Combined RORγ inhibition and anti-PD-L1 blockade reduces neutrophil infiltration and alters myeloid-associated cell populations in orthotopic HCC

To further characterize the effects of RORγ inhibition and anti-PD-L1 blockade on the tumour immune microenvironment, immune and myeloid-associated cell populations were quantified in orthotopic Hep-55.1c tumours following TF-S10, anti-PD-L1, or combination treatment. Combination therapy significantly reduced Ly6G⁺ neutrophil density compared with TF-S10 monotherapy (Figure 6A). Consistent with reduced systemic neutrophil predominance, the peripheral neutrophil-to-lymphocyte ratio was significantly lower in combination-treated mice compared with controls (Figure 6B). The density of RORγ⁺Ly6G⁺ cells was also significantly reduced following combination therapy compared with TF-S10 monotherapy (Figure 6C), indicating decreased accumulation of RORγ-expressing neutrophils within the tumour microenvironment. In contrast, combination treatment significantly increased CD163⁺ cell density compared with controls and both monotherapies (Figure 6D). PD-L1⁺Ly6G⁺ cell density was significantly reduced following combination therapy compared with TF-S10 monotherapy (Figure 6E). Furthermore, RORγ⁺CD163⁺ cell density was significantly increased in combination-treated tumours compared with controls (Figure 6F), suggesting treatment-associated alterations in CD163⁺ cell populations. Collectively, these findings demonstrate that combined RORγ inhibition and PD-L1 blockade reduces tumour-associated neutrophil populations while altering the abundance of CD163⁺ cells, consistent with changes in the cellular composition of the HCC tumour immune microenvironment.

**Fig. 6.**
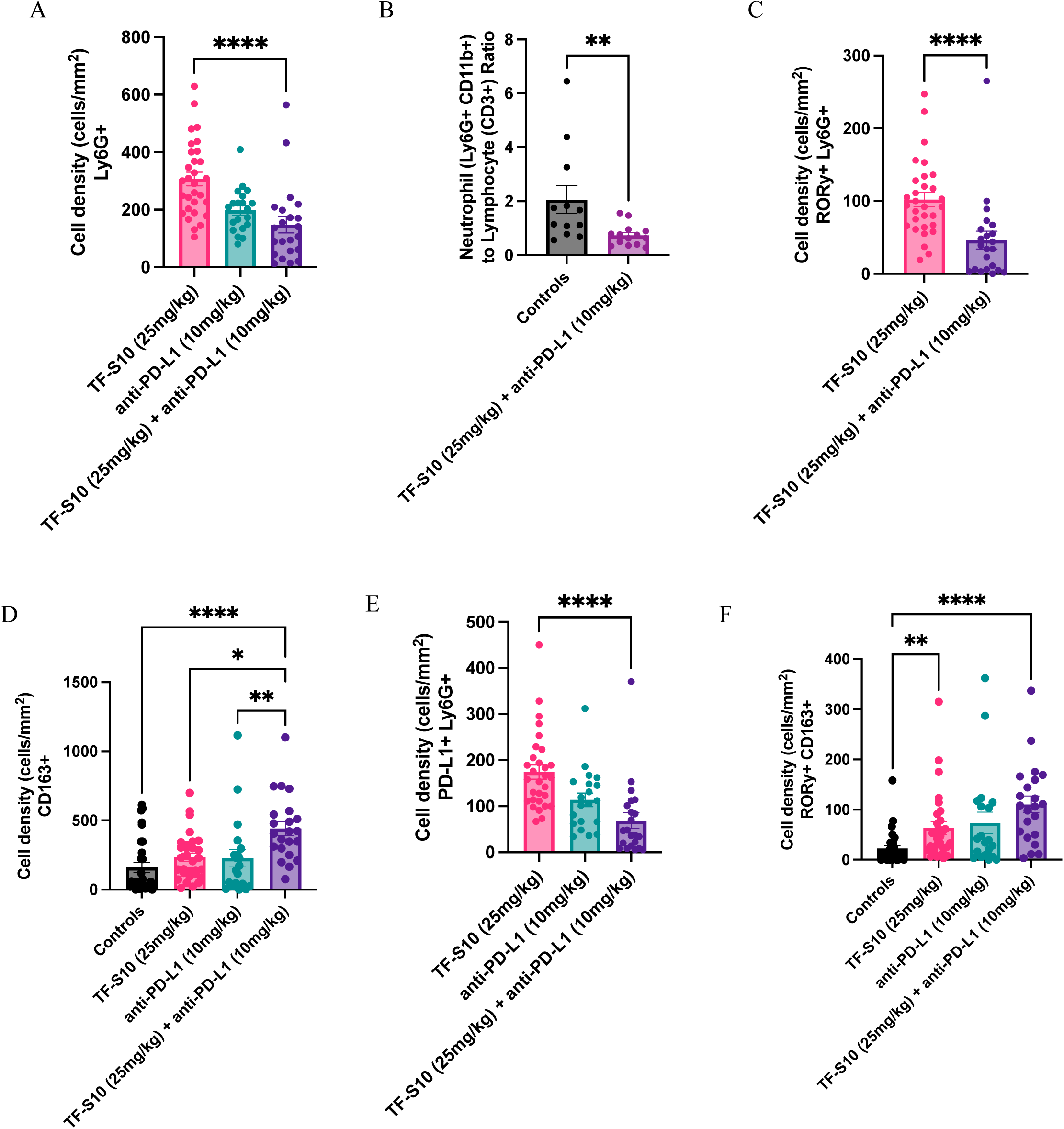
Combined RORγ inhibition and anti-PD-L1 blockade reduces neutrophil-associated inflammation and modulates CD163⁺ cell populations in orthotopic HCC. (A) Quantification of Ly6G⁺ neutrophil density in orthotopic liver tumours from TF-S10 (25 mg/kg), anti-PD-L1 (10 mg/kg), and combination treatment groups. (B) Peripheral neutrophil-to-lymphocyte ratio, calculated as the ratio of Ly6G⁺CD11b⁺ neutrophils to CD3⁺ lymphocytes, in control and combination-treated mice. (C) Quantification of RORγ⁺Ly6G⁺ double-positive cell density in liver tumour sections from TF-S10 and combination treatment groups. (D) Quantification of CD163⁺ cell density in liver tumour sections from control, TF-S10, anti-PD-L1, and combination treatment groups. (E) Quantification of PD-L1⁺Ly6G⁺ double-positive cell density in liver tumour sections from TF-S10, anti-PD-L1, and combination treatment groups. (F) Quantification of RORγ⁺CD163⁺ double-positive cell density in liver tumour sections across all four treatment groups. Cell densities are expressed as cells/mm². Data are presented as mean ± SEM. Statistical significance was assessed using Kruskal–Wallis tests with Dunn’s multiple-comparisons test (A, C–F) or a two-tailed Mann–Whitney U test (B). *P < 0.05, **P < 0.01, ****P < 0.0001.

### RORγ inhibition and anti-PD-L1 treatment reduce pulmonary metastatic burden in orthotopic HCC

To assess whether RORγ inhibition and PD-L1 blockade affect pulmonary metastatic burden, lungs from orthotopic Hep-55.1c tumour-bearing mice were evaluated by hematoxylin and eosin (H&E) staining. Histopathological assessment revealed a shift toward lower pulmonary tumour involvement in treated groups compared with controls, with a greater proportion of TF-S10, anti-PD-L1, and combination-treated mice exhibiting no detectable or low metastatic involvement (Figure 7A). Quantification of the percentage of lung area occupied by tumour demonstrated significantly reduced pulmonary metastatic burden following TF-S10 monotherapy, anti-PD-L1 monotherapy, and combination treatment compared with controls (Figure 7B). However, no significant differences in pulmonary tumour involvement were observed among the treatment groups. Similarly, assessment of pulmonary metastatic foci demonstrated a shift toward fewer metastatic lesions following treatment, with a greater proportion of treated mice exhibiting no detectable foci and fewer mice presenting with moderate or high numbers of metastatic foci (Figure 7C). Representative H&E-stained lung sections further illustrated the differences in pulmonary metastatic involvement across control, TF-S10, anti-PD-L1, and combination-treated groups (Figure 7D). Collectively, these findings demonstrate that RORγ inhibition and PD-L1 blockade, administered individually or in combination, reduce pulmonary metastatic burden in orthotopic HCC, although combination therapy did not provide a statistically significant additional reduction in metastatic area compared with either monotherapy.

**Fig. 7.**
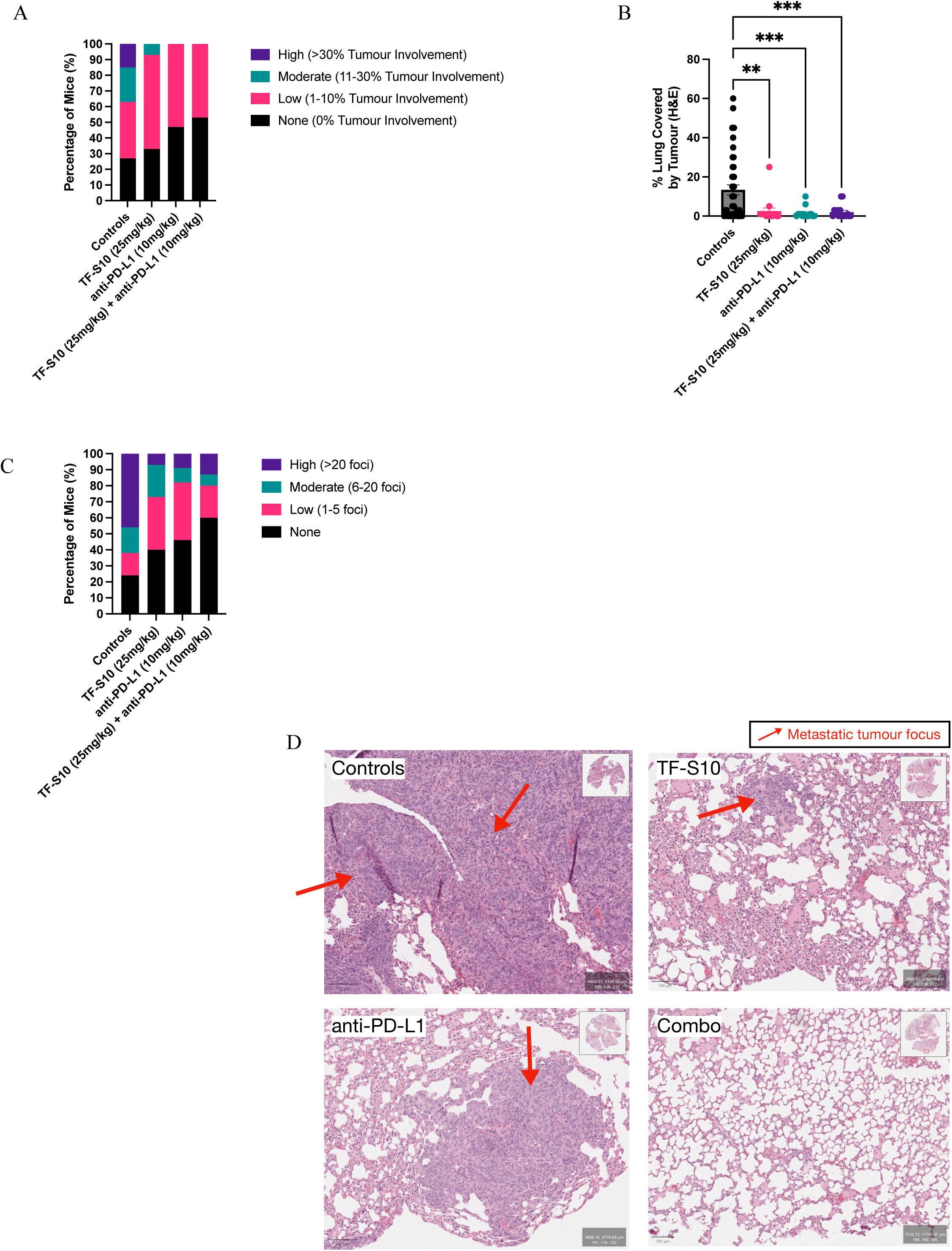
RORγ inhibition and anti-PD-L1 blockade reduce pulmonary metastatic burden in orthotopic HCC. (A) Distribution of mice according to the extent of pulmonary metastatic involvement, categorized as no detectable metastases (0%), low (1–10%), moderate (11–30%), or high (>30%) tumour involvement. (B) Quantification of pulmonary metastatic burden, expressed as the percentage of lung area occupied by tumour, assessed using hematoxylin and eosin (H&E)-stained lung sections from control, TF-S10 (25 mg/kg), anti-PD-L1 (10 mg/kg), and combination treatment groups. (C) Distribution of mice according to the number of pulmonary metastatic foci, categorized as none (0 foci), low (1–5 foci), moderate (6–20 foci), or high (>20 foci). (D) Representative H&E-stained lung sections from control, TF-S10, anti-PD-L1, and combination treatment groups. Red arrows indicate metastatic tumour foci. Insets show low-magnification overviews of the corresponding lung sections. Data in (B) are presented as mean ± SEM. Statistical significance was assessed using Brown–Forsythe and Welch ANOVA followed by Dunnett’s T3 multiple-comparisons test (B). **P < 0.01, ***P < 0.001.

## DISCUSSION

In this study, we identify RORγ as a potential regulator of tumour–immune interactions in HCC, linking its inhibition to reduced neutrophil recruitment, increased tumour-cell PD-L1 expression, and improved tumour control when combined with PD-L1 blockade. Elevated *S100A9* and *CXCL8* (IL-8) expression was associated with reduced overall survival in HCC, consistent with previous reports linking *S100A9* to poor survival and recurrence following HCC resection^15,16^. Tumour-bearing mice also exhibited an increased peripheral neutrophil-to-lymphocyte ratio compared with non-tumour-bearing controls. These findings reinforce the association between neutrophil-related inflammation and adverse HCC outcomes, although neither the transcriptomic associations nor the elevated neutrophil-to-lymphocyte ratio establish a causal relationship with disease progression.

A central finding of this study was that RORγ targeting reduced neutrophil recruitment toward HCC cells. Both *RORC* silencing and pharmacological inhibition with TF-S10 decreased neutrophil migration *in vitro*, while TF-S10 treatment reduced Ly6G⁺ neutrophil infiltration and Ly6G⁺RORγ⁺ cell density in orthotopic liver tumours. HCC-derived chemokines have been implicated in neutrophil recruitment, tumour progression, and metastasis, while tumour-associated neutrophils (TANs) can contribute to immunosuppression and therapeutic resistance^6,8,9,17,18^. Our findings extend these observations by implicating tumour-cell RORγ in the regulation of neutrophil recruitment. However, the soluble mediators responsible for this effect remain unidentified, and direct effects of TF-S10 on host immune cells cannot be excluded. Further investigation is required to establish the underlying mechanisms and determine whether RORγ-mediated changes in neutrophil recruitment influence the development of immune checkpoint inhibitor resistance.

RORγ inhibition also increased PD-L1 expression in HCC cells. Genetic silencing of *RORC* increased *CD274* mRNA expression, while TF-S10 treatment increased PD-L1 expression at both the transcriptional and protein levels. Importantly, the concordant upregulation of CD274 following genetic and pharmacological RORγ inhibition supports an on-target effect of TF-S10, indicating that increased PD-L1 expression is mediated by RORγ inhibition. Although PD-L1 upregulation may facilitate tumour immune evasion, increased expression could also enhance susceptibility to PD-L1-directed blockade. This observation provides a rationale for combining ROR inhibition with anti-PD-L1 therapy, although the extent to which PD-L1 upregulation directly contributes to the improved therapeutic response remains to be established.

In the immunocompetent orthotopic Hep-55.1c model, combined TF-S10 and anti-PD-L1 treatment markedly reduced hepatic tumour burden, as demonstrated by gross tumour volume, histological tumour involvement, and pan-cytokeratin-positive (panCK⁺) tumour-cell density. Combination therapy reduced caliper-measured tumour volume compared with anti-PD-L1 monotherapy and significantly decreased histological tumour involvement compared with either monotherapy. The density of panCK⁺RORγ⁺ tumour cells was also reduced following combination treatment. Although fewer panCK⁺PD-L1⁺ cells were detected *in vivo*, this finding may reflect reduced overall tumour-cell abundance rather than decreased PD-L1 expression at the individual cell level. Quantification of PD-L1 intensity within residual tumour cells would be required to distinguish these possibilities. Furthermore, the absence of longitudinal tumour imaging limits interpretation of treatment kinetics, as endpoint measurements cannot distinguish tumour regression from reduced tumour growth. Future studies incorporating serial imaging would enable direct evaluation of tumour responses over time.

Combination therapy was also associated with changes in the tumour immune microenvironment, including reduced Ly6G⁺, RORγ⁺Ly6G⁺, and PD-L1⁺Ly6G⁺ cell densities. The reduction in peripheral neutrophil-to-lymphocyte ratio further supported a shift in systemic immune-cell composition, although this measurement alone cannot distinguish reduced neutrophil abundance from increased lymphocyte abundance. PD-L1-expressing neutrophils have been implicated in the suppression of adaptive T-cell responses in HCC^19^, suggesting that the observed reduction in these populations may be relevant to the therapeutic response. Nevertheless, functional analyses of neutrophil-mediated immunosuppression and T-cell activity are necessary to establish whether these changes contribute directly to improved antitumour immunity.

In contrast to the reduction in neutrophil-associated populations, combination therapy increased CD163⁺ and RORγ⁺CD163⁺ cell densities, suggesting additional remodelling of myeloid-associated populations. CD163 is commonly expressed by monocytes and macrophages and participates in inflammatory regulation and tissue repair^20–22^, but does not define a uniformly tumour-promoting or immunosuppressive phenotype. Given the substantial reduction in tumour burden, increased CD163⁺ cell density could reflect changes in macrophage recruitment or activation associated with clearance of dying tumour cells and tissue remodelling^23–26^. However, CD163 expression alone cannot establish macrophage functional state or demonstrate efferocytosis, and these populations may encompass overlapping reparative and immunoregulatory phenotypes^27^. Additional phenotypic, functional, and spatial analyses incorporating markers of macrophage identity, activation, immunoregulation, and efferocytosis will be required to clarify their contribution to the therapeutic response^28^.

Beyond hepatic tumour control, RORγ inhibition and PD-L1 blockade were associated with reduced pulmonary metastatic burden, including lower histological lung tumour involvement and a shift toward fewer metastatic foci. All three treatment strategies significantly reduced pulmonary tumour involvement compared with controls; however, combination therapy did not provide a statistically significant additional reduction compared with either monotherapy. These findings are clinically relevant because the lung is a common site of extrahepatic HCC dissemination, and extrahepatic disease generally precludes curative liver transplantation^29,30^. Although transplantation was not directly evaluated, the reduction in hepatic and pulmonary tumour burden supports further investigation of RORγ-targeted approaches for limiting HCC progression and dissemination. Additional limitations include the use of a single immunocompetent orthotopic tumour model and exclusively male mice, which restricts assessment of tumour heterogeneity and potential sex-dependent differences in therapeutic response. Validation in additional HCC models will be important to establish the broader applicability of these findings.

Overall, our findings suggest that RORγ inhibition influences tumour–immune interactions in HCC by reducing neutrophil recruitment while increasing tumour-cell PD-L1 expression. Combined RORγ inhibition and PD-L1 blockade improves hepatic tumour control and alters neutrophil- and CD163-associated cell populations within the tumour microenvironment. These observations provide a rationale for further investigation of RORγ inhibition as a complementary strategy to immune checkpoint blockade, with the potential to improve HCC treatment responses and limit disease progression.

## Acknowledgements

The authors gratefully acknowledge the Flow Cytometry Core Facility at the Research Institute of the McGill University Health Centre (RI-MUHC) for assistance with flow cytometry acquisition and technical support. We thank the RI-MUHC Molecular Imaging Platform for support with immunofluorescence imaging and image acquisition. H&E staining was performed by the RI-MUHC Immunohistochemistry Facility. Multiplex immunofluorescence staining and digital image analysis were performed by the MCIC-Histology Core at BC Cancer in Vancouver; the authors specifically acknowledge Katy Milne, Ming Zi Xu, and Brad H Nelson. This work was supported by funding from the RI-MUHC Endowment Fund, including the Avrith, Cohen, Dolgin, and Guttmann Funds, as well as support from the Montreal General Hospital Foundation and the Courtois Foundation.

## Declaration of generative AI and AI-assisted technologies in the writing process

During the preparation of this manuscript, the authors used ChatGPT (OpenAI) to assist with sentence structure, grammar, and language editing. All content was reviewed and revised by the authors, who take full responsibility for the final manuscript.

## Author Contributions

Sabrina Leo (SL): Conceptualization, Methodology, Investigation, Formal analysis, Data curation, Visualization, Project administration, Writing – original draft, Writing – review & editing; Iqraa Dhoparee-Doomah (IDD): Methodology, Investigation; Sarita Negi (SN): Methodology, Writing – review & editing; Ahmed Fouda (AF): Methodology, Resources; France Bourdeau (FB): Investigation; Gertruda Evaristo (GE): Formal analysis, Writing – review & editing; Steven Paraskevas (SP): Resources, Writing – review & editing; Jonathan Cools-Lartigue (JCL): Conceptualization, Supervision; Jean Tchervenkov (JT): Conceptualization, Methodology, Supervision, Project administration, Writing – review & editing. All authors reviewed and approved the final version of the manuscript.

## Abbreviations

ANOVA: analysis of variance
CTLA-4: cytotoxic T-lymphocyte-associated protein 4
DMEM: Dulbecco’s modified Eagle medium
DMSO: dimethyl sulfoxide
FBS: fetal bovine serum
FPKM-UQ: fragments per kilobase of transcript per million mapped reads upper quartile
HCC: hepatocellular carcinoma
H&E: hematoxylin and eosin
ICI: immune checkpoint inhibitor
IL: interleukin
LIHC: liver hepatocellular carcinoma
NLR: neutrophil-to-lymphocyte ratio
OCT: optimal cutting temperature
PBS: phosphate-buffered saline
PD-1: programmed cell death protein 1
PD-L1: programmed death ligand 1
*RORC*: retinoic acid receptor-related orphan receptor C
RORγ: retinoic acid receptor-related orphan receptor gamma
RORγt: retinoic acid receptor-related orphan receptor gamma t
ROS: reactive oxygen species
RT-qPCR: reverse transcription quantitative polymerase chain reaction
SEM: standard error of the mean
siRNA: small interfering RNA
TAN: tumour-associated neutrophil
TCGA: The Cancer Genome Atlas
Treg: regulatory T cell

---

**Supplementary Table 1.**
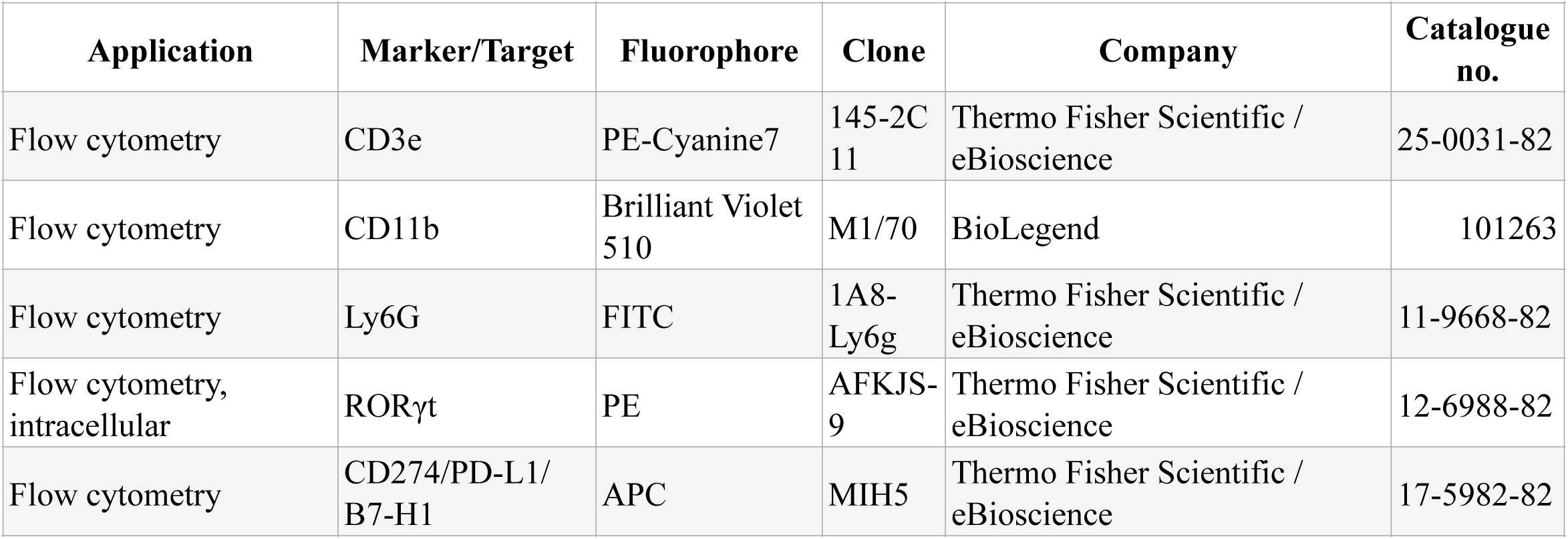
Antibodies used for flow cytometric analysis. Antibodies used to assess immune-cell populations and PD-L1 expression by flow cytometry. Surface staining included CD3e, CD11b, Ly6G, and CD274/PD-L1, while intracellular staining included RORγt. Antibody clone, fluorophore, vendor, and catalogue information are provided.

